# Localization of AP2α2, TRPV1 and PIEZO2 to the Large Dense Core Vesicles of Human Dorsal Root Ganglion Neurons

**DOI:** 10.1101/2025.03.31.646357

**Authors:** Ojasi Joshi, Aubrey Cooper, Rasheen Powell, Molly K. Martin, Raider Rodriguez, Joseph B. Kuechle, Arin Bhattacharjee

## Abstract

Dorsal Root Ganglia (DRG) consist of both peptidergic and non-peptidergic nociceptive neurons. CGRP, an inflammatory neuropeptide, is a classical marker of peptidergic nociceptors and CGRP is stored within the large dense core vesicles (LDCVs) of these neurons. In addition to storing large peptide neurotransmitters, LDCVs might also serve to transport key membrane proteins to the peripheral terminals. This immunohistochemical study investigated the localization of different membrane proteins to the LDCVs of human DRG neurons. Previously validated antibodies against the endocytotic subunit AP2α2, the heat-activated channel TRPV1 and the mechanosensitive channel PIEZO2 were used in conjunction with an antibody against CGRP on sections of intact human DRG isolated from de-identified human subjects. Immunohistochemical studies were also performed on human synovial tissue to examine peripheral terminals. High magnification confocal microscopy was used to determine the co-localization signal of these membrane proteins with CGRP. We observed a strong co-localization of AP2α2 with the CGRP containing LDCVs signifying its role in membrane recycling. Moreover, we also observed a strong colocalization of TRPV1 and PIEZO2 with CGRP suggesting that LDCV release controls the trafficking of these channels to the membrane. It is likely that during injury, bulk exocytosis of CGRP will concomitantly increase the surface expression of TRPV1 and PIEZO2 channels enhancing the responsiveness of these neurons to painful stimuli. This model suggests that neurons that co-localize TRPV1 and PIEZO2 to CGRP containing LDCVs are likely silent nociceptors.

## 1 Introduction

Regulated exocytosis from neurons involves at least two types of secretory vesicles: small secretory vesicles (SSVs) and large dense core vesicles (LDCVs). SSVs contain classical neurotransmitters and undergo repeated cycles of exo-endocytosis in nerve terminals, constituting synaptic transmission. However, peptide neurotransmitters must be packaged into LDCVs, which, when viewed under electron microscopy, have a characteristic dense core (Coulter, 1988; De Camilli & Jahn, 1990). In rodent neurons, SSVs have a diameter of 40-50 nm, while LDCVs range from 100-500 nm in diameter (Lin et al., 2022). Clathrin-mediated endocytosis is an essential component of vesicular secretion, enabling membrane retrieval after exocytosis (Jung & Haucke, 2007).

Neurons utilize the multimeric adaptor protein complex 2 (AP2) to recruit clathrin and facilitate endocytosis (AP2-CME) after exocytosis (Santos, Li, & Voglmaier, 2009). The AP-2 adaptor complex comprises four subunits: α, β2, μ2, and σ2: (Robinson, 2004; Traub, 1997, 2003). Notably, there are two genes that encode the α subunit: α*1* and α*2*. The α1 subunit is localized to synaptic compartments, while the α2 subunit is localized to extrasynaptic compartments (Ball, Hunt, & Robinson, 1995). Previously, we demonstrated that AP2α2 subunit is preferentially expressed in calcitonin gene-related peptide (CGRP)^+^ mouse and human dorsal root ganglion (DRG) neurons (Powell et al., 2021).

Nociceptive DRG neurons are classified as either peptidergic or non-peptidergic. CGRP and substance P (SP) are two common markers of peptidergic nociceptors. These neuropeptides are packaged into LDCVs within the trans-Golgi network of the somata of neurons and subsequently transported to the spinal cord or peripheral terminals, where they are eventually secreted. SP and CGRP are categorized as being neuroinflammatory peptides because during an injury, they are released en masse to coordinate inflammation. However, these peptides also have numerous interoceptive functions, including regulation of bone metabolism and local vasodilation. For instance, CGRP is a potent vasodilator that regulates core body temperature. CGRP-containing nociceptors express the heat-sensitive transient receptor potential vanilloid 1 (TrpV1) channel (Hwang, Oh, & Valtschanoff, 2005; Premkumar & Sikand, 2008) and it was shown that TrpV1 channel antagonists affected core body temperature through the attenuated release of CGRP(Yue, Yuan, Braz, Basbaum, & Julius, 2022). Therefore, the release of these peptides during non-pain conditions serves various homeostatic functions.

Despite ongoing research, the roles of peptidergic neurons and their contributions to pain signaling during inflammation remain areas of active investigation. One class of peptidergic dorsal root ganglion (DRG) neurons are the silent nociceptors. These neurons are typically unresponsive to mechanical stimulation but become responsive during injury and inflammation. During inflammatory pain, normally mechanically insensitive nociceptors become sensitized to both mechanical and thermal stimuli, leading to the development of mechanical and thermal hyperalgesia (Dubin & Patapoutian, 2010). It is estimated that silent nociceptors constitute approximately 30% of all C-fiber afferents in viscera and joints and about 15–20% in human skin, while they appear to be less abundant in mouse skin (Nees et al., 2023). There is evidence suggesting that the mechanically sensitive channel Piezo2 is responsible for transducing mechanical stimuli in silent nociceptors (Obeidat et al., 2023; Prato et al., 2017). However, the molecular mechanisms underlying the transition of silent nociceptors to the sensitized state remain unclear. One potential hypothesis is that simultaneous membrane trafficking of TrpV1 and Piezo2 occurs during inflammation.

In this study we immunohistochemically characterized AP2α2, TRPV1 and PIEZO2 in human DRG neurons. Human DRG neurons are two to ten times larger than mouse DRG neurons (Haberberger, Barry, Dominguez, & Matusica, 2019). To our surprise, CGRP containing LDCV were visible using immunofluorescence and high magnification. This enabled us to examine the co-localization of membrane associated proteins at LDCV offering a putative model as to how silent nociceptors “wake up”.

## 2 Materials and Methods

### 2.1 Animals

Experiments and procedures conducted on C57Bl/6 mice were approved by the University at Buffalo Institutional Animal Care and Use Committee (IACUC) in accordance with the guidelines set forth by the National Institutes of Health. Adult mice were purchased from Envigo (Indianapolis, IN).

### 2.2 Antibody Characterization

Table 1 provides all the primary antibody identifiers and dilutions/concentrations used in the study. The CGRP antibody (Abcam, Cambridge MA #ab81887), has been cited in over 150 publications. The AP2α2 antibody (Abcam #ab220065 was previously characterized using genetic knockdown of AP2α2 (Chen et al., 2018; Powell et al., 2021). The TrpV1 antibody (Invitrogen, Carlsbad, CA, #PA1-29770) was characterized by immunohistochemistry against rat DRG neurons (manufacturer’s data sheet) and been cited in multiple publications (Ding et al., 2023; La Montanara et al., 2020). The Piezo2 antibody (Novus Biologicals, Centennial, CO #NBP1-78624 is the most cited antibody on Piezo2 and was validated using Piezo2 floxed mice(Fang et al., 2021).

**Table 1:**
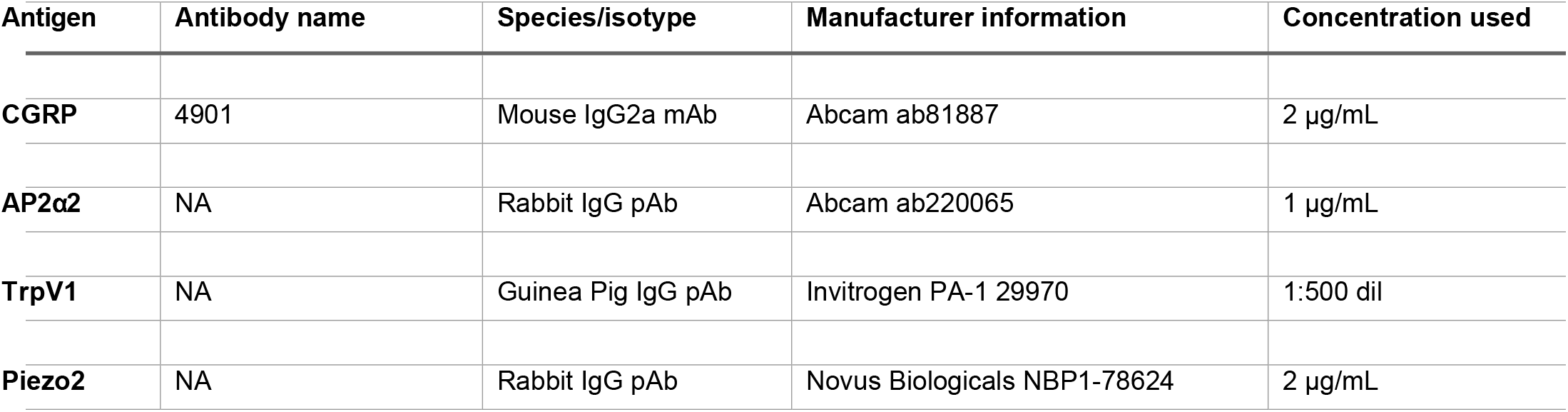
Primary Antibodies.

### 2.3 Immunohistochemistry Human Dorsal Root Ganglia

Human DRG were acquired from the University at Buffalo Anatomical Gift Program. The donors were 74 years old female, 83 years old female and 77 years old male with no medical history. No identifying information was shared with the researchers. DRG were initially preserved in formaldehyde, rehydrated, sequentially, in decreasing ratios of phosphate buffered saline (PBS) to water: 24 hours in 50% PBS then 24 hours in 30% PBS. Following rehydration, the DRG were cryoprotected in 30% sucrose at 4°C and submerged in Tissue Freezing Media, TFM™ (Electron Microscopy Sciences, Hatfield, PA) and frozen using dry-ice chilled 2-methylbutane. Once the resulting blocks were thoroughly frozen, they were placed into a -80°C freezer for 48 hours. Slices for staining were made on cryostat at 16 microns for DRGs (mouse and human). Cryosections were taken and mounted onto charged Superfrost microscope slides.

Sections were first washed 3 times with PBS, then incubated overnight in blocking media (10% Normal Goat Serum (Abcam, Cambridge, MA), 3% Bovine Serum Albumin (Sigma, St. Louis, MO) and 0.025% Triton X-100 (Sigma) in PBS. Slides were then incubated in primary antibodies for 24h at 4°C. Slides were then incubated with the secondary antibodies (goat anti-mouse 555 1:1000, donkey anti-rabbit 647, goat anti-guinea pig 488 (all from Invitrogen/ThermoFisher Scientific) for 24h at 4°C. The following day, the slides were rinsed 4 times with PBS. then mounted on slides using Vectashield Antifade Mounting Media with 4,6-diamidino-2-phenylindole dihydrochloride (DAPI; Life Technologies/ ThermoFisher Scientific, Waltham, MA).

All slides were allowed 24h to set at 4°C, before imaging. All lower magnification images were acquired using a Leica DMi8 inverted fluorescent microscope equipped with a sCMOS Leica camera (Leica, Wetzlar, Germany) and connected to a Hewlett Packard Z4 G4 Workstation loaded with THUNDER enabled LAS X imaging software. Higher magnification images were acquired using a Leica TCS SP8 Confocal Microscope under oil immersion. Optimal antibody concentration and exposure time, for each antibody and tissue type, was determined at initial tissue imaging. The same exposure time was used for each tissue type. After exposure, min/max range was selected for optimal contrast and to minimize background. All images were analysed using a separate workstation that was loaded with the LAS X imaging software. Images were further modified (e.g. addition of scale bars) using Adobe Illustrator.

### 2.4 Immunohistochemistry Human Synovium

Synovial tissue was donated by a consenting female patient (Age 61) who underwent knee joint replacement surgery at Buffalo General Medical Center (Buffalo, NY). The tissue donation was consistent with IRB guidelines according to protocol as approved: Orthopaedic Tissue Procurement University at Buffalo IRB study 1729. No other identifying information was shared with the researchers. Samples were sectioned at 20 microns and placed on microscope slides (Fisherbrand, Pittsburgh, PA). Sections were washed 3 times with a wash buffer (0.4% Triton X-100 in PBS), then incubated overnight in blocking solution (10% normal goat serum, 3% bovine serum albumin, and 0.4% Triton X-100 in PBS). The next day, slides were incubated in primary antibodies overnight at 4°C. The following day, slides were incubated in secondary antibodies overnight at 4°C. Afterwards, slides were washed twice with wash buffer and mounted using Vectashield antifade mounting medium with DAPI (VectorLabs, Newark, CA).

### 2.5 Immunohistochemistry Mouse Dorsal Root Ganglia

Mice were anesthetized with a high dose of sodium pentobarbital and perfused transcardially with 4% paraformaldehyde as previously described(Gage, Kipke, & Shain, 2012). Lumbar DRGs were removed from the fixed mice and postfixed overnight. Tissue was then cryoprotected in PBS containing 30% sucrose overnight. Mouse DRG were sectioned and stained in a similar manner described above for the human DRG using the same antibody concentrations. DRG Slices for staining were made at 16 microns.

## 3 Results

### 3.1 AP2α2 and TrpV1 localize to CGRP^+^ DRG neurons

CGRP is expressed in a subset of mouse DRG neurons(Ruscheweyh, Forsthuber, Schoffnegger, & Sandkuhler, 2007). We recapitulated the preferential expression of Ap2α2 (Figure 1a) and TrpV1 channels (Figure1b) to CGRP-containing DRG neurons. For CGRP expression in human DRG neurons, at lower magnification, punctate CGRP immunolabeling was already observable and AP2α2 co-localized to this CGRP puncta (Figure 1c).

**Figure 1.**
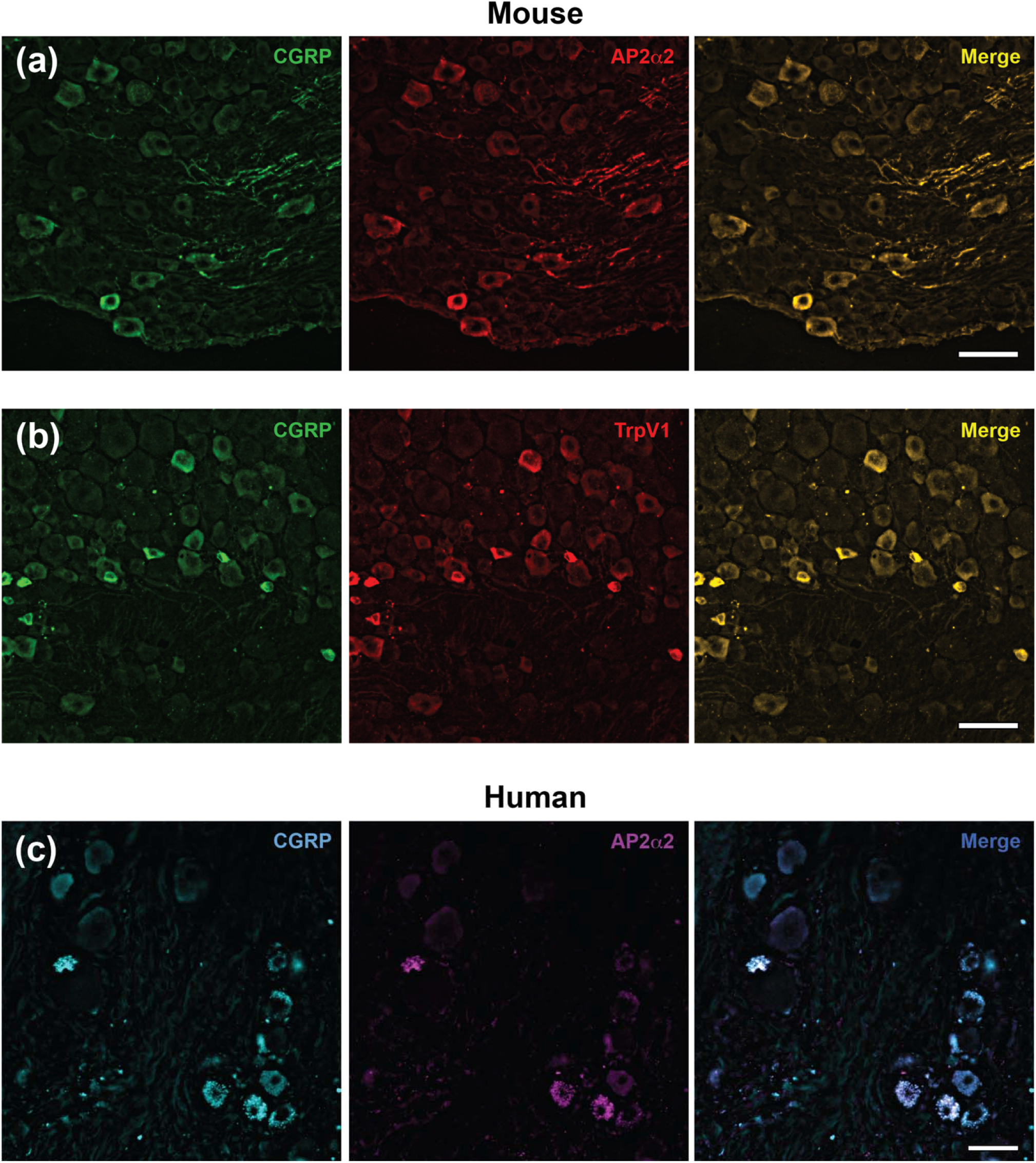
AP2α2 and TRPV1 are localized to CGRP+ neurons a) AP2α2 is expressed in mouse CGRP^+^ DRG neurons replicating prior work (Powell et al., Nat Comm 2021 (Scale bar 50μm). TRPV1 channels are expressed in in mouse CGRP^+^ DRG neurons replicating prior work (Deves et al. PNAS 2014)(Scale bar 50μm). c) AP2α2 is expressed in human CGRP^+^ DRG neurons replicating prior work (Powell et al., Nat Comm 2021 (Scale bar 190μm).

### 3.2 High magnification immunofluorescence microscopy reveal AP2α2 immunolocalization to CGRP-containing LDCVs in human DRG neurons

Observing CGRP puncta in human DRGs at lower magnification prompted us to visualize the immunofluorescence at 40X magnification (Figure 2a) and at 100X magnification (Figure 2b). We found robust AP2α2 immunolabeling localized to CGRP-containing vesicles in 3 independent subjects.

**Figure 2.**
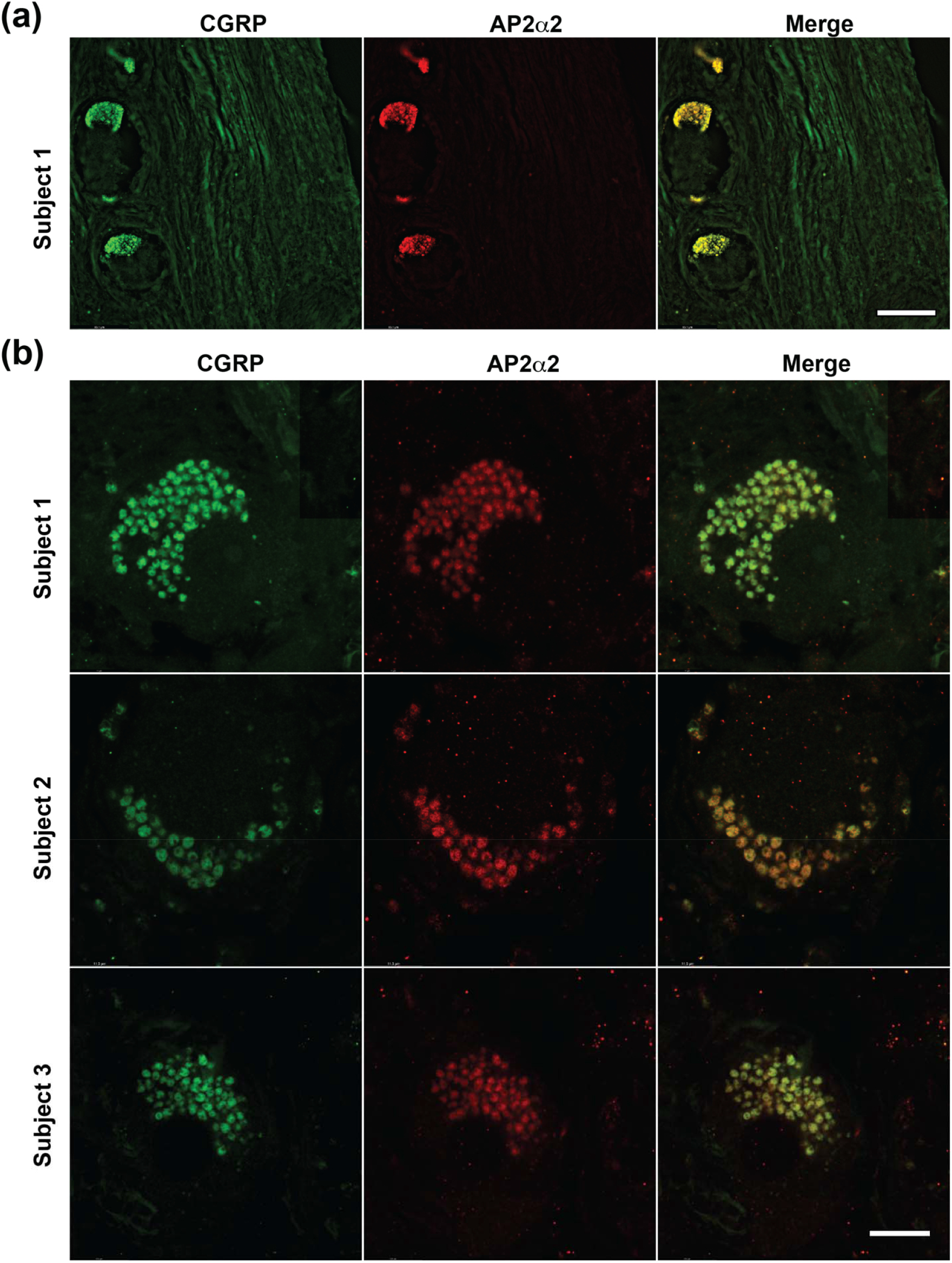
Under higher magnification AP2α2 colocalizes with CGRP containing LDCVs a) Images of human DRG neurons taken at 40X magnification (Scale bar 190μm). b) Images of human DRG neurons taken under 100X magnification in 3 different subjects. (Scale bar 50μm).

### 3.3 High magnification immunofluorescence microscopy reveal TRPV1 immunolocalization to CGRP-containing LDCVs in human DRG neurons

Prior studies have shown that TrpV1 channels localize to LDCVs in mouse DRG neurons(Devesa et al., 2014). We accordingly probed for TRPV1 in human DRG neurons and found overlapping immunoreactivity with CGRP containing LDCVs under high magnification (Figure 3a). In a different human subject, we performed a CGRP, AP2α2 and TRPV1 co-localization study finding all three proteins localizing to LDCVs (Figure 3b). Using three-dimensional reconstruction under high magnification, it’s clear the immunolabeling occurred at spherical LDCVs within the neuron.

**Figure 3.**
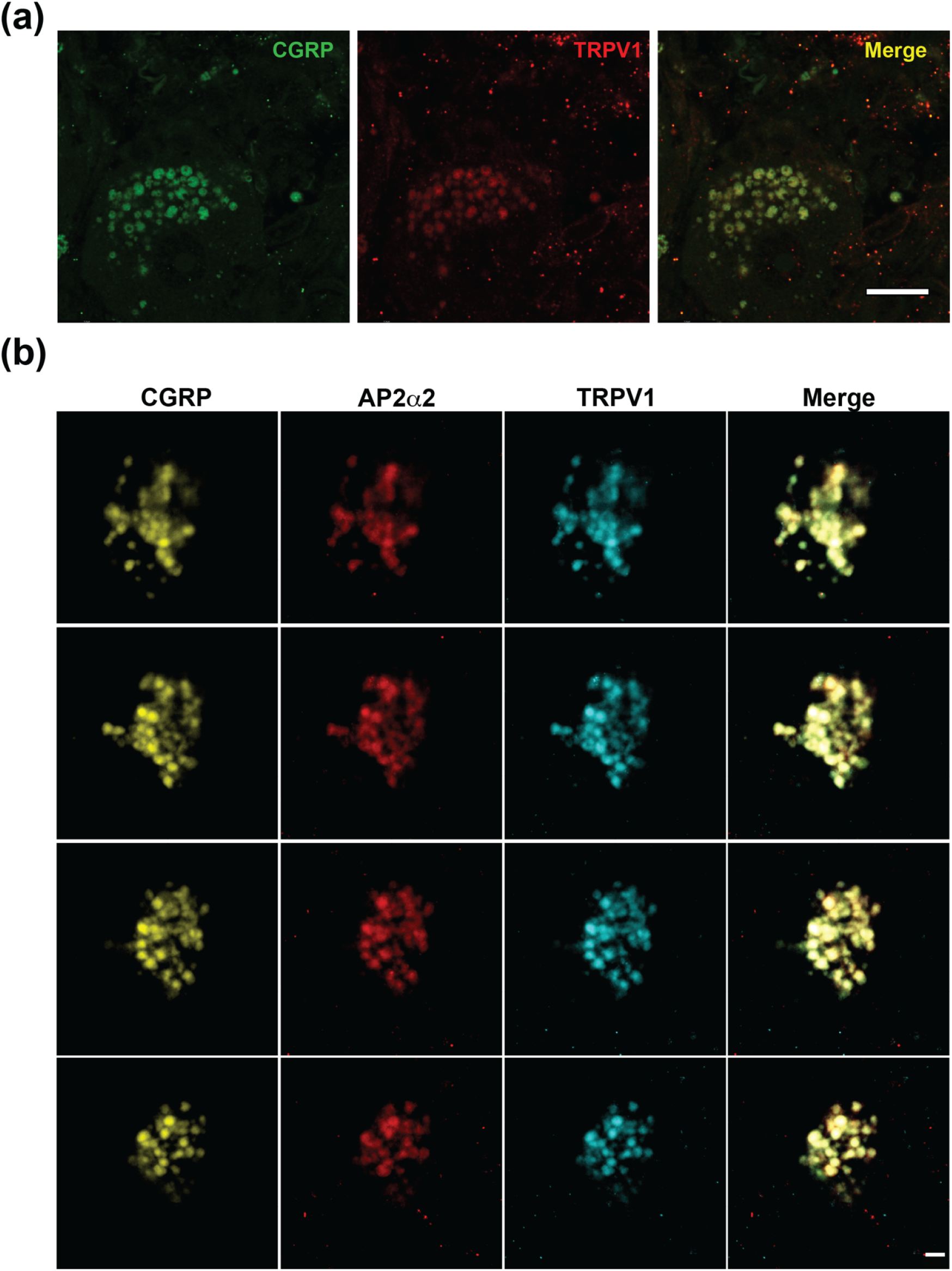
Under higher magnification AP2α2 and TRPV1 colocalize with CGRP containing LDCVs a) Images of human DRG neurons taken under 100X magnification (Scale bar 50μm). b) 3D reconstruction of a human DRG stained for AP2α2, TRPV1 and CGRP at 100X magnification (Scale bar 5 μm)

### 3.4 Piezo 2 localization in human DRG neurons

Unlike the selective expression of TrpV1 channels in a subset of mouse DRG neurons (Figure 1b), Piezo2 channels exhibit widespread distribution in mouse DRG neurons (Shin et al., 2021). In human DRG neurons, we similary observed widespread distribution of PIEZO2 (Figure 4). Expression appeared robust at the membrane, however in CGRP^+^ DRG neurons, we were observed intracellular PIEZO2 immunolabeling with co-localization to CGRP in 3 independent subjects (Figure 4).

**Figure 4.**
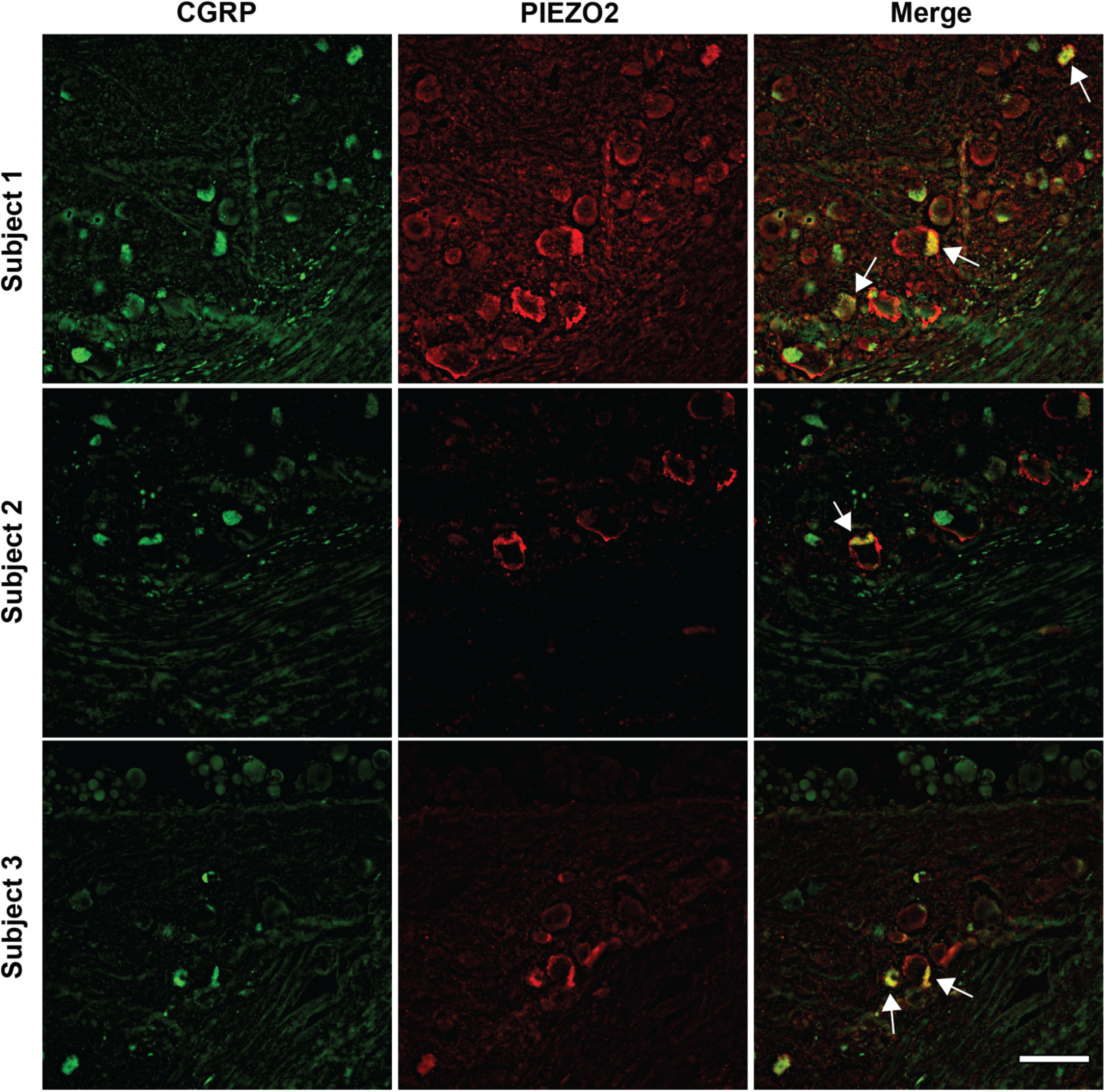
PIEZO2 shows overlapping and non-overlapping localization in CGRP+ neurons CGRP and Piezo2 immunolocalization in human DRG neurons under 40X magnification. Piezo2 immunolabeling exhibited both overlapping and non-overlapping localization with CGRP. Arrows indicate areas of co-localization Scale bar 100 μm

### 3.5 High magnification immunofluorescence microscopy reveal PIEZO2 immunolocalization to CGRP-containing LDCVs in human DRG neurons

With increasing magnification, we were able to resolve PIEZO2 immunoreactivity at CGRP-containing LDCVs (Figure 5). At 63X both membrane and LDCV localization of PIEZO2 were visible (Figure 5a) and at 100X, the localization of PIEZO2 to LDCVs was like that of AP2a2 and TRPV1 (Figures 2&3),

**Figure 5.**
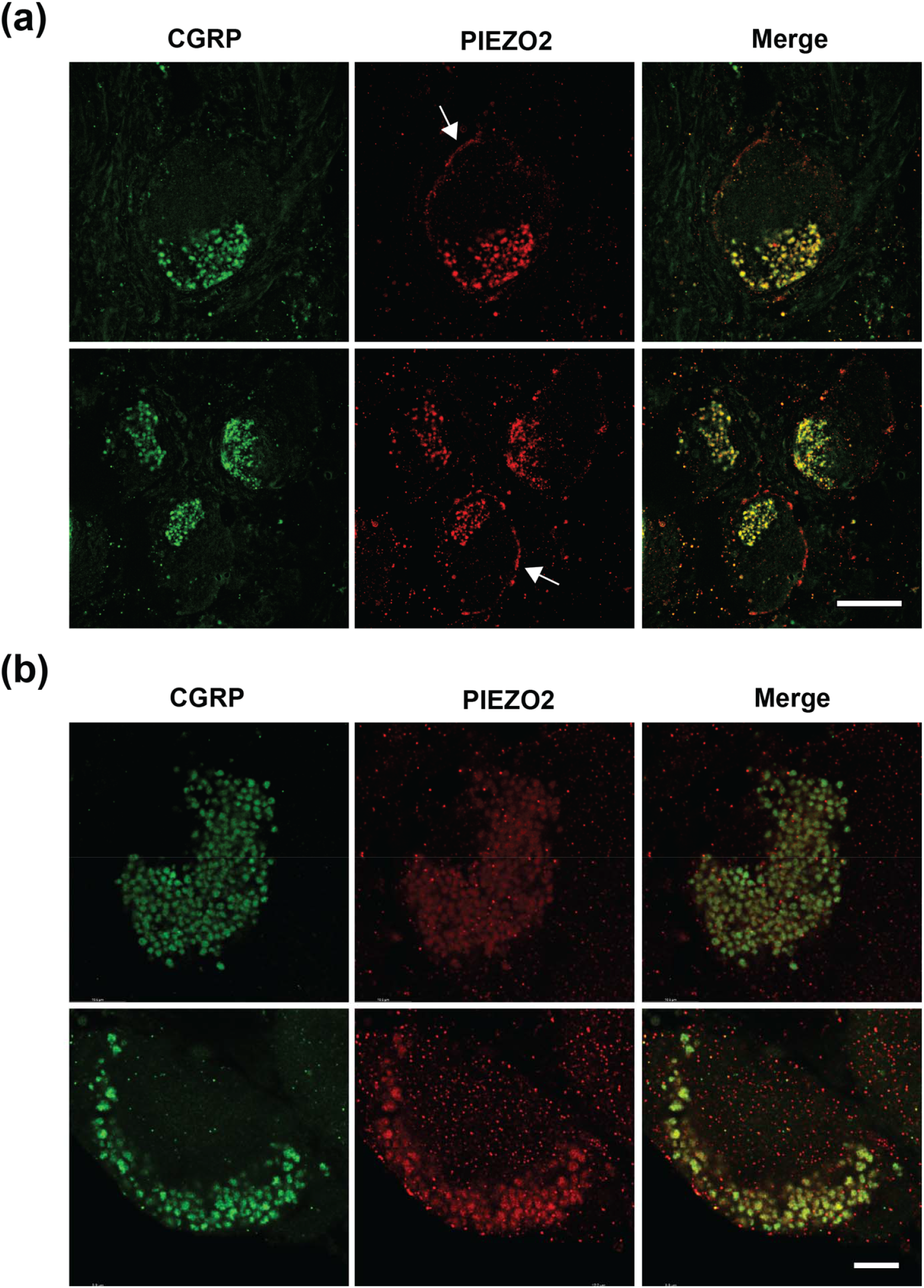
Under higher magnification PIEZO2 colocalizes with CGRP containing LDCVs a) Images of human DRG neurons taken at 63X magnification (Scale bar 110μm). Arrows indicate membrane localization of a pool of PIEZO2 channels b) Images of human DRG neurons taken under 100X magnification (Scale bar 10μm).

### 3.6 High magnification immunofluorescence microscopy reveal TRPV1 and PIEZO2 co-immunolocalization in human DRG neurons

We repeated high magnification immunofluorescence microscopy imaging with PIEZO2 and TRPV1 labels in 3 independent subjects. We observed robust co-localization of channels in presumed LDCVs.

### 3.7 Trafficking of LDCVs containing CGRP and AP2α2 to the periphery

It has been estimated that 75% of knee afferents in rat DRG express CGRP (Staton, Wilson, Bountra, Chessell, & Day, 2007) and in back-labeling knee afferents experiments no IB4 binding was observed(Fernihough, Gentry, Bevan, & Winter, 2005). To understand whether CGRP-containing LDCVs transport AP2α2 to the periphery we subsequently performed immunolabeling studies on isolated human synovial tissue donated from a patient that had undergone knee arthroplasty. First, we confirmed the presence of unmyelinated fibers in the synovial tissue using peripherin labeling (Figure 7a). Peripherin^+^ fibers expressed both CGRP and AP2α2(Figure 7a). Under high magnification, like what was observed at DRG somata, we observed spherical, overlapping CGRP and AP2α2 immunoreactivity (Figure 7b). We believe that these were CGRP-containing LDCVs that were being transported to the periphery.

### 3.8 Trafficking of LDCVs containing CGRP, TRPV1 and PIEZO2 to the periphery

TrkA receptors bind to nerve growth factor (NGF) and NGF is known to sensitize TrpV1 channels(Zhang, Huang, & McNaughton, 2005). Recent work identified a subset of nociceptors that express both TRKA and PIEZO2 transcripts in human lumbar DRG neurons and Piezo2 channels mediate mechanical sensitivity in experimental arthritis in mice(Obeidat et al., 2023). Under high magnification we again observed spherical CGRP containing vesicles that also contained TRPV1 and PIEZO2 immunoreactivity, demonstrating that these channels are transported to peripheral terminals via LDCV vesicular transport.

### 3.9 Model of LCDV meditated transport of AP2α2, TRPV1 and PIEZO2

This schematic summarizes how three proteins, AP2α2, TRPV1 and PIEZO2, are transported from the soma to peripheral terminals via CGRP-containing LDCVs. Under injury conditions, where there is increased CGRP release, TRPV1 and PIEZO2 membrane expression is increased causing increased sensitivity to painful stimuli. AP2α2 as part of the AP2 complex is important for membrane retrieval after LDCV exocytosis.

## 4 Discussion

The term “large dense core vesicles” was coined by electron microscopy examination of rodent specimens. In our study, we utilized human DRG and synovial tissue to immunofluorescently visualize CGRP-containing large dense core vesicles. As with larger-sized neurons, LDCVs in human DRG neurons appear 5-10 times larger than LDCVs in rodents. Using human DRGs enabled us to ascertain whether other proteins are localized to LDCVs, and we discovered that AP2α2, TRPV1, and PIEZO2 all localize to LDCVs.

### 4.1 The AP2 complex alpha2 gene is associated with LDCVs in human DRG neurons

LDCVs can undergo both kiss-and-run exocytosis and full-collapse exocytosis(Artalejo, Elhamdani, & Palfrey, 1998). For full-collapse exocytosis, classical endocytosis is crucial for maintaining the supply of vesicles needed for continued secretion(Artalejo et al., 1998). For non-peptidergic DRG neurons, endocytosis is required for synaptic transmission of SSVs in the dorsal horn. It is plausible to assume that for peptidergic neurons, there is additional endocytotic pressure to maintain membrane homeostasis following LDCV exocytosis at both central and peripheral terminals. It’s not surprising that the AP2α2 subunit is preferentially expressed in IB4^-^ peptidergic nociceptors(Powell et al., 2021), however, we observed here that the AP2α2 directly localized to CGRP-containing LDCVs at DRG somata and in peripheral terminals of synovial tissue (Figures 2&7). This suggests that AP2-CME is closely associated with the exocytosis of LDCV in DRG neurons. It remains to be determined whether the other components of the AP2 complex, namely β2, μ2, and σ2, are also localized to LDCVs.

### 4.2 Multiple proteins localize to LDCV

In our previous research, we determined that Slick sodium-activated potassium channels (Kcnt2) were expressed in CGRP-containing mouse DRG neurons and Slick channels localized to LDCVs (Tomasello, Hurley, Wrabetz, & Bhattacharjee, 2017).

Furthermore, we demonstrated that neuronal stimulation leading to CGRP release resulted in the translocation of Slick channels to the neuronal membrane (Tomasello et al., 2017). Similarly, TrpV1 channels localized to LDCV in mouse neurons also mobilized to the surface membrane following stimulation(Devesa et al., 2014). In a liquid chromatography-mass spectrometry study of rat dorsal horns, up to 298 proteins were found associated with LDCV membranes. Ion channels, G-protein-coupled receptors and signaling molecules comprised this group of proteins. The precise mechanism by which proteins are sorted into LDCVs remains unclear. It is unknown whether these proteins contain an LDCV localization sequence that directs them to LDCVs or if targeting to LDCVs is a mere default pathway for these proteins. For AP2α2 and TRPV1 we saw almost exclusive localization of proteins to the LDCVs (Figures 2&3), while for PIEZO2 both LDCV and membrane localization was observed (Figures 4&5). PIEZO2 channels are extensively spliced with various isoforms(Szczot et al., 2017) and this may determine the differential targeting of channels to LDCVs or to the plasma membrane.

Nevertheless, neuropeptide synthesis, packaging into LDCVs, and transport to the periphery are continually ongoing for peptidergic neurons. It is plausible to associate the transport of various proteins with LDCV transport to the periphery as an energy-saving mechanism. Again, the sorting and localization of these proteins to LDCVs in peptidergic neurons might reflect a simple default pathway. This remains to be further explored.

### 4.3 Neurons that exhibit LDCV localization of bothTRPV1 and PIEZO2 may function as silent nociceptors

Silent nociceptors are typically unresponsive to painful stimuli under normal conditions. but become activated only in response to inflammation or tissue injury. PIEZO2 is required for mechanosensitivity in silent nociceptors and NGF sensitizes silent, but not other nociceptors to mechanical stimuli(Prato et al., 2017). There is a higher percentage of TRPV1/TRKA1-positive neurons in human DRG neurons compared to mouse DRG neurons (Rostock, Schrenk-Siemens, Pohle, & Siemens, 2018). Therefore, silent nociceptors are likely to express both TRPV1 and PIEZO2 channels particularly in human DRG neurons. The simultaneous recruitment of both channels to the membrane during injury or inflammation explains how silent nociceptors “wake up” (i.e. cause thermal and mechanical hypersensitivity). Molecularly, how this simultaneous recruitment occurs has not been well defined. From our observational studies on LDCV localization of both channels (Figures 6&8) we present a model (Figure 9) on the unsilencing of these nociceptors. Injury causes increased CGRP release, a phenomenon first described many decades ago(Donnerer & Stein, 1992). The full collapse of CGRP containing LDCVs would allow the translocation of many TRPV1 and PIEZO2 channels to the neuronal membrane sensitizing these neurons to thermal, acidic pH and mechanical stimuli. In other words, increased LDCV exocytosis is the mechanism by which silent nociceptors awaken. From a therapeutic point of view blocking CGRP release might be a therapeutic approach to treat pain. Unfortunately, it was shown that treatment of trigeminal ganglia cultures with therapeutic concentrations of botulinum toxin type A did not affect the amount of CGRP released from these neurons(Durham, Cady, & Cady, 2004). We previously showed that locally inhibiting nociceptor endocytosis with an AP2 complex decoy peptide caused CGRP retention in the superficial layers of the epidermis(Powell et al., 2021). It could be that by blocking endocytosis in this way, prevented full maturation of LDCVs and blocked the recruitment of necessary synaptic proteins required for exocytosis. Nonetheless, further studies of the molecular machinery required for LDCV exocytosis in DRG neurons is warranted for the targeting of silent nociceptors.

**Figure 6.**
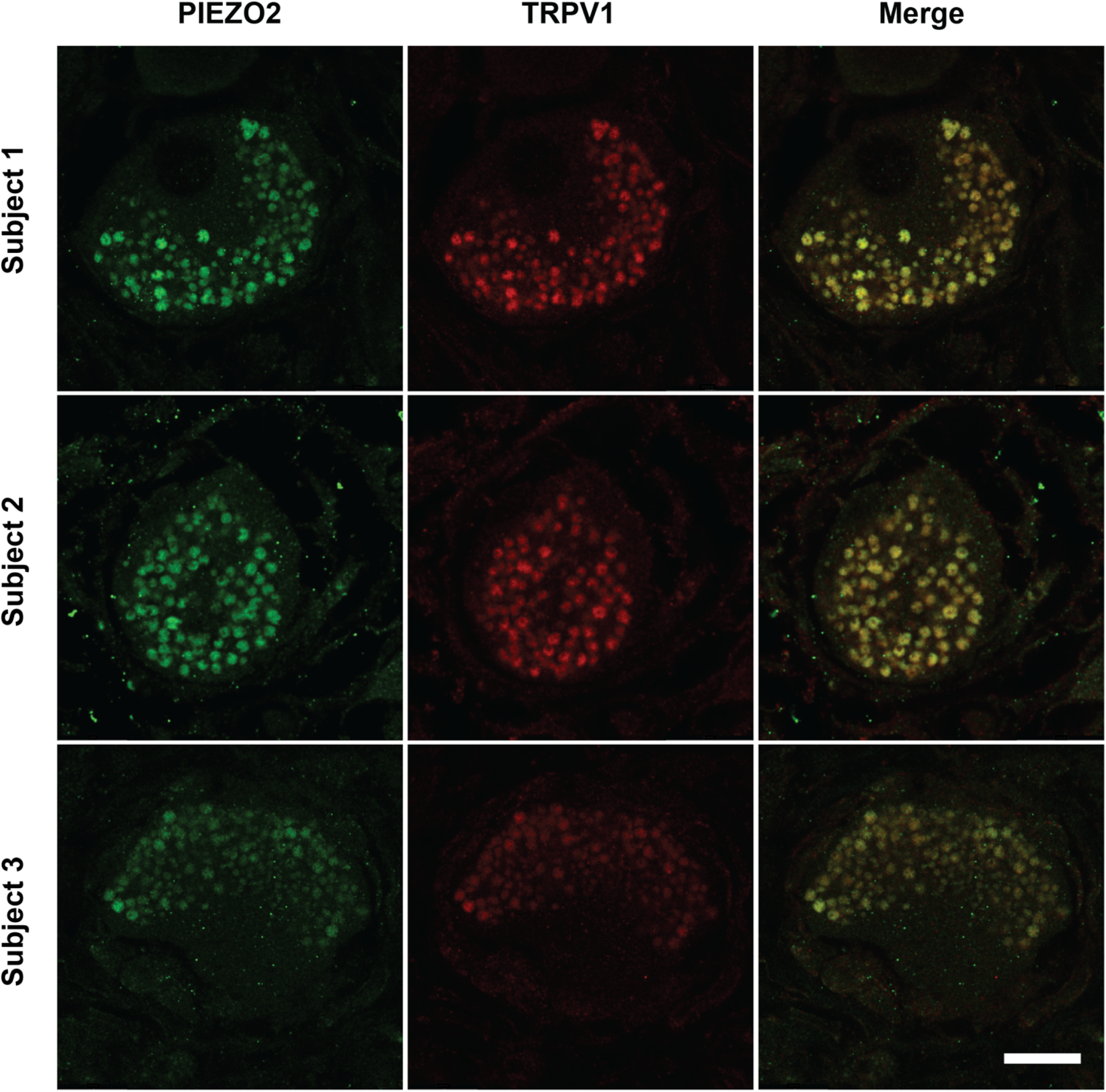
TRPV1 and PIEZO2 colocalization in presumed LDCVs. Images of human DRG neurons taken under 100X magnification in 3 different subjects. (Scale bar 20μm).

**Figure 7.**
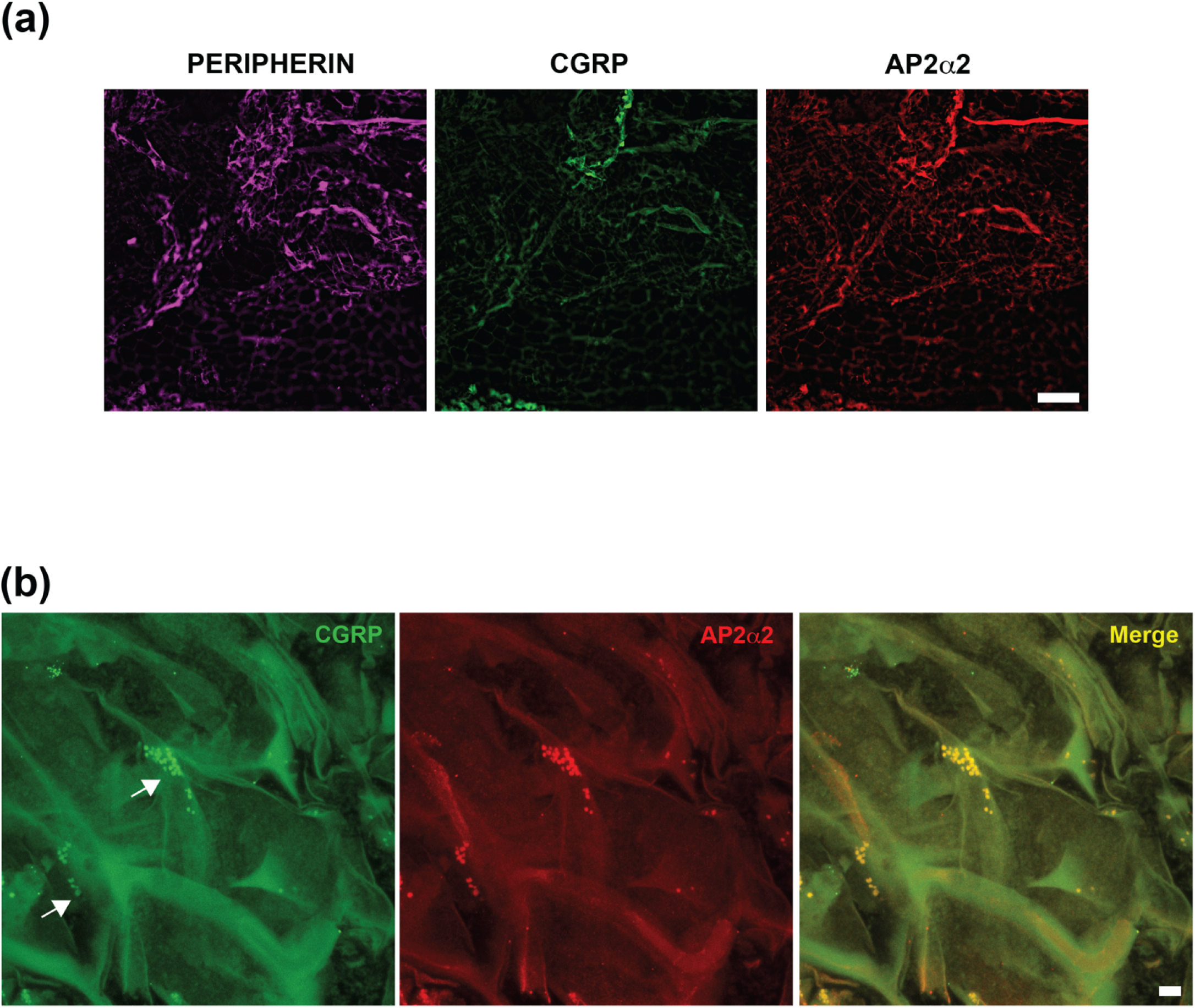
AP2α2 is transported to the periphery by CGRP-containing LDCVs. a) Images of unmyelinated nociceptors in human synovial tissues. Peripherin is a marker for unmyelinated neurons and these nociceptive fibers contain CGRP and Ap2α2. (Scale bar 100μm) b) Images of human synovial tissue taken under 100X magnification. Arrows indicate CGRP containing LDCVs. Ap2α2 immunolabeling exhibited overlapping immunoreactivity with CGRP. (Scale bar 5μm)

**Figure 8.**
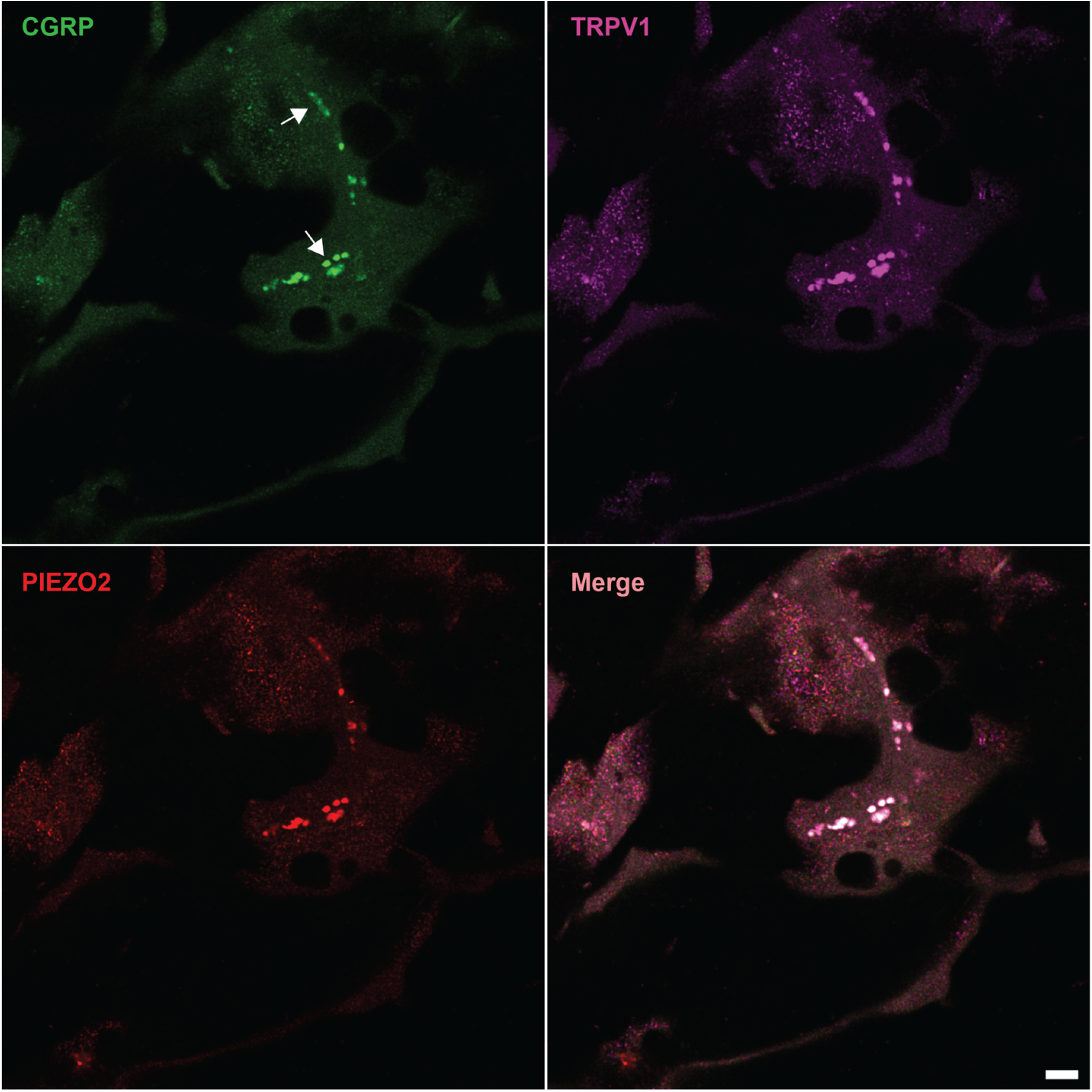
TRPV1 and PIEZO2 are transported to the periphery by CGRP-containing LDCVs. Images of human synovial tissue taken under 100X magnification. Arrows indicate CGRP containing LDCVs. Both TRPV1 and PIEZO2 immunolabeling exhibited overlapping immunoreactivity with CGRP. (Scale bar 5μm)

**Figure 9.**
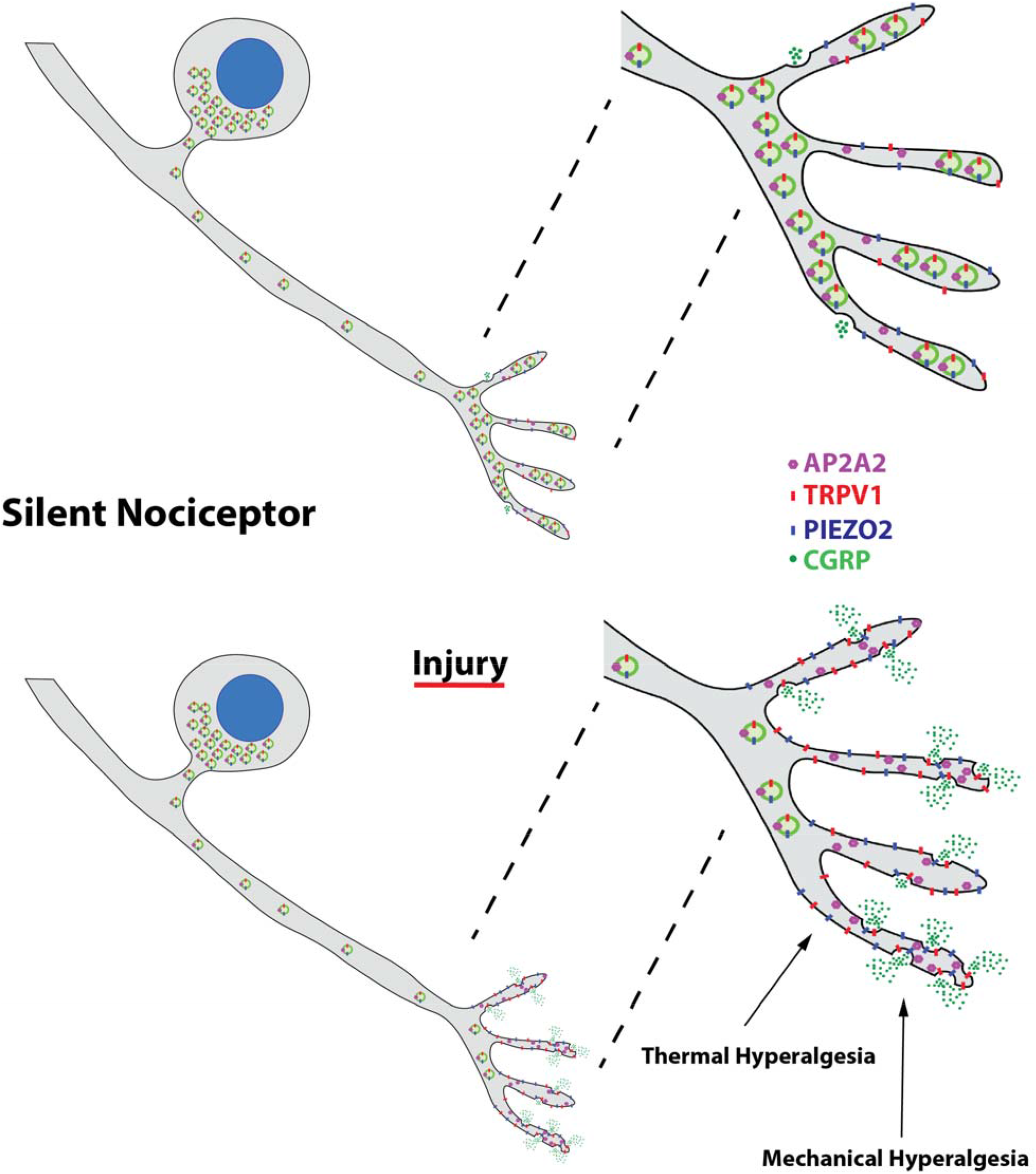
Schematic model for LDCV mediated transport of AP2α2, TRPV1 and PIEZO2.

## Author Contributions

AB, OJ, and RP conceived the idea for the project and co-wrote the manuscript. OJ performed all immunohistochemistry experiments on human DRG and analyzed the data associated with human DRG. AC performed all immunohistochemistry experiments on human synovial tissue and analyzed the data associated with human synovial tissue. AC and MKM isolated DRG from human cadavers. RR processed synovial tissue. JBK isolated human synovial tissue.

## Acknowledgments

This work was supported by NS128543. The authors would like to thank to thank Dr. Wade Sigurdson of the University at Buffalo Confocal Microscopy Core Facility for his assistance with the confocal imaging.

